# A tRNA modification balances carbon and nitrogen metabolism by regulating phosphate homeostasis, to couple metabolism to cell cycle progression

**DOI:** 10.1101/507707

**Authors:** Ritu Gupta, Adhish S. Walvekar, Shun Liang, Zeenat Rashida, Premal Shah, Sunil Laxman

## Abstract

Cells must appropriately sense and integrate multiple metabolic resources to commit to proliferation. Here, we report that cells regulate carbon and nitrogen metabolic homeostasis through tRNA U_34_-thiolation. Despite amino acid sufficiency, tRNA-thiolation deficient cells appear amino acid starved. In these cells, carbon flux towards nucleotide synthesis decreases, and trehalose synthesis increases, resulting in a starvation-like metabolic signature. Thiolation mutants have only minor translation defects. However, these cells exhibit strongly decreased expression of phosphate homeostasis genes, resulting in an effectively phosphate-limited state. Reduced phosphate enforces a metabolic switch, where glucose-6-phosphate is routed towards storage carbohydrates. Notably, trehalose synthesis, which releases phosphate and thereby restores phosphate availability, is central to this metabolic rewiring. Thus, cells use thiolated tRNAs to perceive amino acid sufficiency, and balance carbon and amino acid metabolic flux to maintain metabolic homeostasis, by controlling phosphate availability. These results further biochemically explain how phosphate availability determines a switch to a ‘starvation-state’.

## Introduction

Cells utilize multiple mechanisms to sense available nutrients, and appropriately alter their internal metabolic state. Such nutrient-sensing systems assess internal resources, relay this information to interconnected biochemical networks, and control global responses that collectively reset the metabolic state of the cell, thereby determining eventual cell fate outcomes (Jeong *et al.*, 2000; Förster *et al.*, 2003; Zaman *et al.*, 2008; Broach, 2012; Cai and Tu, 2012; Ljungdahl and Daignan-Fornier, 2012) However, much remains unknown about how cells sense and integrate information from multiple nutrient inputs, to coordinately regulate the metabolic state of the cell and commit to different fates.

In this context, the metabolic state of the cell is also closely coupled with protein translation. Protein synthesis is enormously energy consuming, and therefore must be carefully regulated in tune with nutrient availability (Warner, 2001). Generally, overall translational capacity and output increases during growth and proliferation (Jorgensen *et al.*, 2004), and decreases during nutrient limitation (Wullschleger, Loewith and Hall, 2006). Signaling processes that regulate translational outputs (such as the TORC1 and PKA pathways) are well studied (Wullschleger, Loewith and Hall, 2006; Zaman *et al.*, 2008; Broach, 2012; González and Hall, 2017). Notwithstanding this, little is known about how core components of the translation machinery might directly control metabolic outputs, and thus couple metabolic states with physiological cellular outcomes.

tRNAs are core components of the translation machinery, and are extensively modified post-transcriptionally (Björk *et al.*, 1987; Phizicky and Hopper, 2010). Some tRNA modifications are required for tRNA folding, stability, or the accuracy and efficiency of translation (Phizicky and Hopper, 2010). However, the roles of many of these highly conserved modifications remain unclear. One such modification is a thiolation of uridine residue present at the wobble-anticodon (U_34_) position of specifically glu-, gln- and lys-tRNAs (s^2^U_34_) (Gustilo, Vendeix and Agris, 2008; Phizicky and Hopper, 2010). In yeast, this is mediated by a group of six enzymes-Nfs1, Tum1, Uba4, Urm1, Ncs2 and Ncs6, which are evolutionarily conserved (Nakai, Nakai and Hayashi, 2008; Leidel *et al.*, 2009; Noma, Sakaguchi and Suzuki, 2009). These enzymes incorporate a thiol group derived directly from an amino acid (cysteine), and replace the oxygen present at the 2-position of U_34_ with sulfur (Schmitz *et al.*, 2008; Leidel *et al.*, 2009; Noma, Sakaguchi and Suzuki, 2009). Surprisingly, these thiolated tRNAs appear to have a relatively minor role in general translation, as seen in multiple studies (Rezgui *et al.*, 2013; Zinshteyn and Gilbert, 2013; Klassen *et al.*, 2016; Chou *et al.*, 2017) with modest roles in enhancing the efficiency of wobble base codon-anticodon pairing (Yarian *et al.*, 2002; Rezgui *et al.*, 2013).

Contrastingly, tRNA thiolation appears to directly alter cellular metabolism, but this connection has remained largely unexplored. Thiolated tRNAs are required to maintain metabolic cycles in yeast (Laxman *et al.*, 2013). Further, the amounts of thiolated tRNAs reflect the intracellular availability of sulfur-containing amino acids (cysteine and methionine) (Laxman *et al.*, 2013). Several studies have observed that a loss of this modification alters amino acid homeostasis, inducing the amino-acid starvation regulator Gcn4 (Laxman *et al.*, 2013; Zinshteyn and Gilbert, 2013; Nedialkova and Leidel, 2015). Additionally, studies have found that the loss of tRNA thiolation results in hypersensitivity to oxidative agents, and the TORC1 inhibitor rapamycin (Fichtner *et al.*, 2003; Goehring, 2003; Goehring, Rivers and Sprague, 2003; Laxman and Tu, 2011; Scheidt *et al.*, 2014), all supporting a role for thiolated tRNAs in maintaining metabolic homeostasis. These disparate studies hint that a core function of this tRNA modification may be to sense amino acid availability (primarily methionine/cysteine), and somehow regulate the metabolic state of the cell to appropriately commit to growth. Yet, how thiolated tRNAs regulate metabolism, and the extent to which this may control cellular outcomes remains entirely unaddressed.

In this study, by directly analyzing different metabolic outputs, we identify the metabolic nodes that are altered in tRNA thiolation deficient cells. We find that tRNA thiolation regulates central carbon and nitrogen (amino acid) metabolic outputs, by controlling flux towards storage carbohydrates. In tRNA thiolation deficient cells, overall metabolic homeostasis is altered, with flux diverted away from the pentose-phosphate pathway/nucleotide synthesis axis, and towards storage carbohydrates trehalose and glycogen. This thereby alters cellular commitments towards growth and cell cycle progression. Counter-intuitively, we discover that this metabolic-state switch in cells lacking tRNA thiolation is achieved by down-regulating a distant metabolic arm of phosphate homeostasis. We biochemically elucidate how regulating phosphate balance can couple amino acid and carbon utilization towards or away from nucleotide synthesis, and identify trehalose synthesis as the pivotal control point for this metabolic switch. Through these findings we show how tRNA thiol-modifications couple amino acid sensing with overall metabolic homeostasis. We further present a general biochemical explanation for how inorganic phosphate homeostasis regulates commitments to different arms of carbon and nitrogen metabolism, thereby determining how cells commit to a ‘growth’ or ‘starvation’ state.

## Results

### Amino acid and nucleotide metabolism are decoupled in tRNA thiolation deficient cells

Earlier studies observed an increased expression of amino acid biosynthetic genes, and an activation of the amino acid starvation responsive transcription factor Gcn4, in cells lacking tRNA thiolation (Laxman *et al.*, 2013; Zinshteyn and Gilbert, 2013; Nedialkova and Leidel, 2015). These studies therefore suggested that tRNA thiolation-deficient cells were amino-acid starved. We investigated this surmise, by directly measuring free intracellular amino acids in wild-type (WT) and tRNA thiolation mutant cells (*uba4*Δ and *ncs2*Δ). Note: these two independent pathway mutants were chosen, to avoid misinterpretations coming from possible roles independent of tRNA thiolation. We used prototrophic yeast strain grown in synthetic minimal medium without supplemented amino acids, in order to minimize any possible confounding roles of supplemented amino acids/nucleotides in the medium. We compared relative amounts of amino acids in WT and thiolation mutants by using quantitative, targeted LC-MS/MS approaches. Here, we carefully normalized samples by total cell number (based on optical density at 600 nm), as well as biomass. Unexpectedly, we observed a substantial increase in the intracellular pools of amino acids in thiolation mutants (Figure 1A). This shows that the thiolation deficient cells are not amino acid starved, but contrarily accumulate amino acids. We next correlated these actual amino acid amounts with the abundance of Gcn4. Gcn4 is the major amino acid starvation responsive transcription factor, and is induced upon amino acid starvation (Hinnebusch, 1984, 2005) (Figure 1B). We measured Gcn4 protein amounts in WT and thiolation deficient cells, and contrarily observed increased Gcn4 protein in thiolation mutants (Figure 1C). Further, *GCN4* translation was correspondingly higher in thiolation mutants (Figure S1A) (as also seen earlier in (Zinshteyn and Gilbert, 2013; Nedialkova and Leidel, 2015), and this increased *GCN4* translation in the thiolation mutants was Gcn2- and eIF2α phosphorylation-dependent (Figure S1B and Figure S1C). These observations comparing actual amino acid amounts in cells with the activity of Gcn4 therefore present a striking paradox. As canonically understood, Gcn4 is induced upon amino acid starvation, and Gcn4 translation and protein decrease when intracellular amino acid amounts are restored (Hinnebusch, 1984, 2005). Contrastingly, in the results observed here, despite the high amino acid amounts in the tRNA thiolation mutants, the Gcn2-Gcn4 pathway remains induced. We therefore concluded that the metabolic node regulated by tRNA thiolation, resulting in an apparent amino acid starvation signature, cannot be at the level of amino acid biosynthesis and availability.

**Figure 1:**
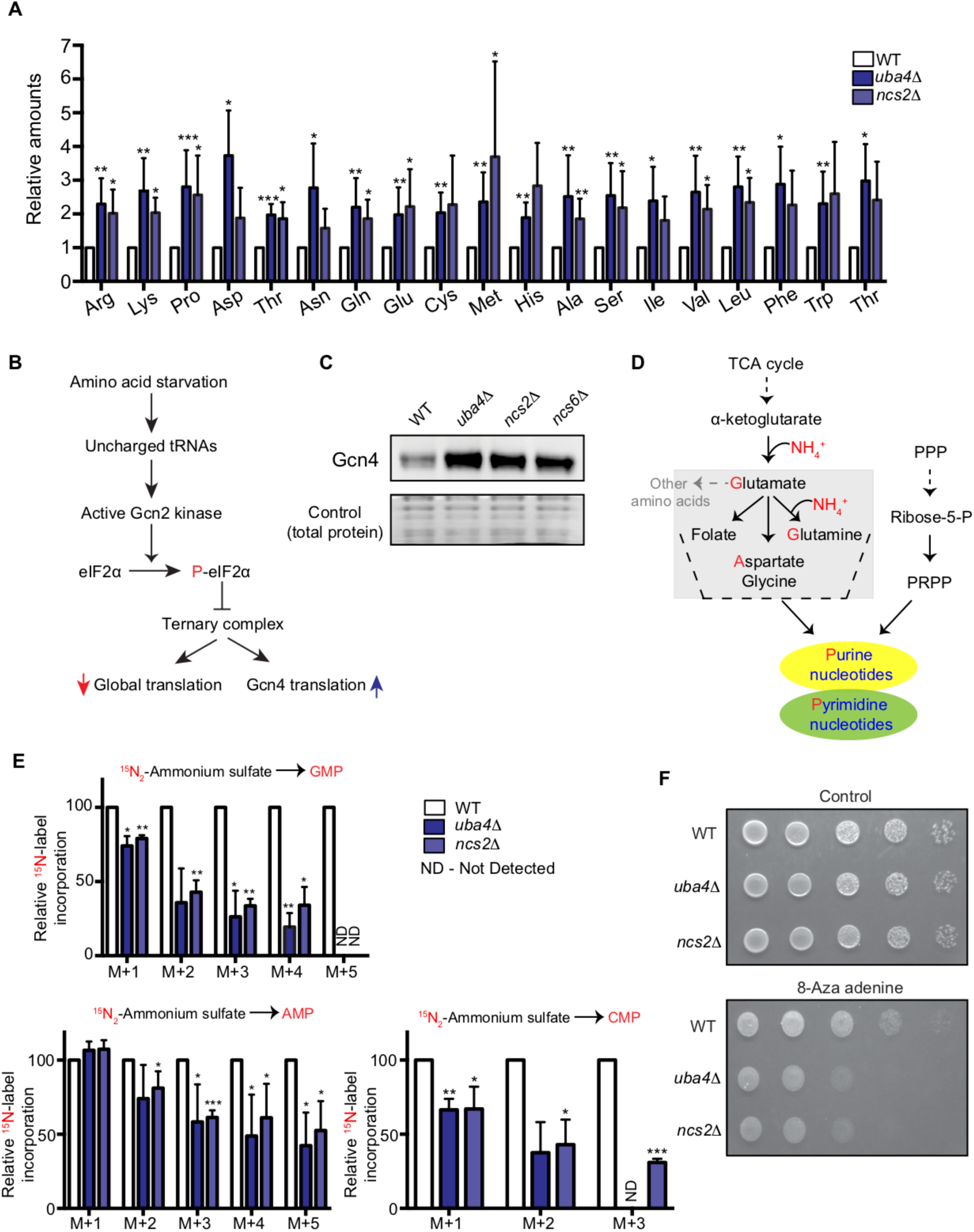
Amino acid and nucleotide metabolism are decoupled in tRNA thiolation deficient cells. (A) Intracellular pools of amino acids are increased in tRNA thiolation mutants. Steady-state amino acid amounts were measured in wild-type (WT) and tRNA thiolation mutant cells (*uba4*Δ and *ncs2*Δ) grown in minimal media by targeted liquid chromatography/mass spectrometry (LC-MS/MS). Data are displayed as means ± SD, n>=3. *p<0.05, **p<0.01, ***p<0.001. (B) A schematic representation illustrating the induction of Gcn4 translational upon amino acid starvation, as mediated by the Gcn2 kinase, and phosphorylation of the eIF2α initiation factor. (C) Gcn4 protein is increased in tRNA thiolation mutants. Western blots indicating Gcn4 protein levels (Gcn4 tagged with HA epitope at the endogenous locus) in WT and tRNA thiolation mutant cells (*uba4*Δ, *ncs2*Δ and *ncs6*Δ) grown in minimal media, as detected using an anti-HA antibody. A representative blot obtained from three biological replicates (n=3) is shown. Also see figures S1A, S1B and S1C. (D) A schematic representation of *de novo* nucleotide (purine and pyrimidine) biosynthesis as made from its precursors-amino acids, folates and PRPP (5-Phopsphoribosyl-1-Pyrophosphate). (E) Nucleotide synthesis is decreased in tRNA thiolation mutants. WT and tRNA thiolation mutant cells (*uba4*Δ and *ncs2*Δ) grown in minimal media were pulse-labelled with ^15^N_2_-labelled ammonium sulfate for 90 minutes to measure newly synthesized nucleotides (GMP, AMP and CMP) using targeted LC-MS/MS. Label incorporation in tRNA thiolation mutant cells relative to WT is plotted, where label incorporation in WT was set to 100%. The incorporation of ^15^N atoms from ^15^N_2_-labelled ammonium sulfate into nucleotides is represented as M+n, where n is the number of ^15^N-labelled atoms. Data are displayed as means ± SD, n=3 for AMP and CMP, n=2 for GMP. *p<0.05, **p<0.01, ***p<0.001. Also see Figures S2A, S2B and S2D. (F) tRNA thiolation mutants exhibit increased sensitivity to 8-azaadenine. WT and tRNA thiolation mutant cells (*uba4*Δ and *ncs2*Δ) grown in minimal media were spotted on minimal media agar plates containing 300 µg/ml 8-Aza adenine. Also see Figures S2C.

We therefore considered the possible metabolic outcomes of amino acid utilization, and hypothesized that an alteration in amino acid utilization could be a source of this metabolic rewiring. In particular, amino acids are the sole nitrogen donors for *de novo* nucleotide synthesis (Figure 1D). Since amino acids accumulated in thiolation mutants, we explored the possibility that this was due to reduced *de novo* nucleotide synthesis. For this, we adopted a metabolic flux based approach to unambiguously estimate new nucleotide synthesis (Walvekar, Rashida, *et al.*, 2018). In such an approach, unlabelled or ^15^N-labelled ammonium sulfate can be provided as a sole nitrogen source, and label incorporation via glutamine and aspartate into newly formed nucleotides can be measured. Notably, the incorporation of ^15^N-label into nucleotides (GMP, AMP and CMP) decreased in thiolation mutants relative to WT cells (Figure 1E), indicating reduced flux towards nucleotide synthesis. Consistent with this, we also observed decreased steady-state levels of nucleotides in thiolation mutants (Figure S2A). Notably, the ^15^N-label incorporation into amino acids (aspartate and glutamine) themselves were not affected in thiolation mutants (Figure S2B), ruling out amino acid synthesis defects. We further addressed this pharmacologically, using a purine-analog, 8-azaadenine, which acts as a pseudo-feedback inhibitor of nucleotide biosynthesis. Consistent with the decreased nucleotide levels observed, thiolation mutants exhibited increased sensitivity to 8-azaadenine (Figure 1F, and S2C). Collectively, these data show that the loss of tRNA thiolation decreases nucleotide biosynthesis, with a corresponding accumulation of amino acids. Indeed, nucleotide limitation is an established mechanism to induce Gcn4 (Rolfes and Hinnebusch, 1993). We also asked if the decreased nucleotide synthesis was due to reduced expression of nucleotide biosynthetic genes. We observed that the expression of candidate genes in this pathway were increased in thiolation mutants (Figure S2D), diminishing the possibility of reduced nucleotide biosynthetic capacity as a reason for decreased nucleotides. This is further explored later in this manuscript. Indeed, increased mRNA levels of nucleotide biosynthetic genes observed in thiolation mutants may be due to feedback upregulation in response to reduced nucleotides, which is a well-established phenomenon (Davis and Ares, 2006; Kwapisz *et al.*, 2008).

Collectively, despite increased intracellular pools of amino acids, tRNA thiolation deficient cells exhibit signatures of amino acid starvation, including decreased nucleotide biosynthesis. These data show that tRNA thiolation is important for cells to appropriately balance amino acid utilization with nucleotide synthesis.

### Carbon flux is routed towards storage carbohydrates in thiolation mutants

Despite amino acids being non-limiting in thiolation deficient cells, flux towards nucleotide synthesis was decreased. This observation was in itself puzzling, and the reason was not obvious. We therefore asked if carbon metabolism was instead rewired in the thiolation mutants. Our reasoning was as follows: while amino acids are the sole nitrogen donors for nucleotide synthesis, the carbon backbone for nucleotides is derived from central carbon metabolism (Figure 2A). We reasoned that since the decreased nucleotide synthesis was not due to amino acid limitation, this could instead be due to a metabolic shift where carbon flux is routed away from nucleotide synthesis. Carbon derived from glucose is converted to glucose-6-phosphate, and then is typically directed towards glycolysis and the pentose phosphate pathway (PPP), which provides precursors (ribose sugars) for nucleotide synthesis (Figure 2A) (Boyle, 2005). However, if glucose-6-phosphate is instead diverted towards storage carbohydrates trehalose and glycogen (Figure 2A), this will result in reduced flux into the PPP, and thereby decreased nucleotide synthesis. To assess if this might happen in tRNA thiolation mutants, we pulsed [U-^13^C_6_]-labelled glucose to growing WT or thiolation deficient cells, and measured label incorporation into different central carbon metabolites, with nucleotides as the end-point readout. We observed significantly decreased carbon label incorporation towards new nucleotide synthesis (GMP and AMP) in the thiolation mutants (Figure 2B and S3A). This result is also consistent with the decreased nucleotide synthesis based on amino acid derived nitrogen assimilation, observed earlier (Figure 1E). Note: in this time-frame it is technically difficult to estimate ^13^C-glucose incorporation into glycolytic and pentose phosphate pathway intermediates (2/3-phosphoglycerate, 6-phosphogluconate, ribose-5-P/ribulose-5-P, sedoheptulose-7-P), since the carbon-label incorporation rapidly saturates (Van Heerden *et al.*, 2014). Therefore, as expected, amounts of these labelled metabolites were unchanged in WT and thiolation mutant cells (Figure S3B).

**Figure 2:**
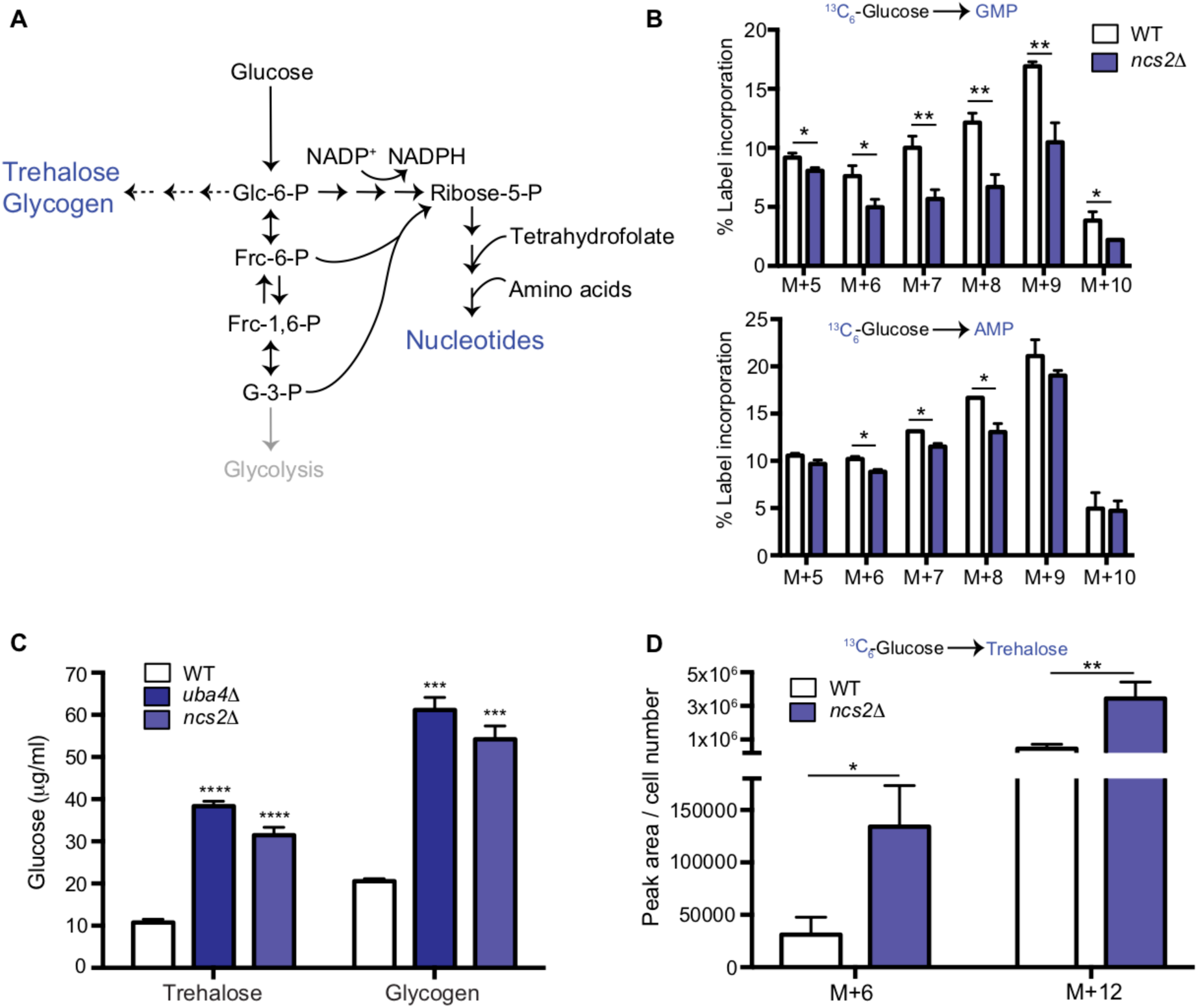
Carbon flux is routed towards storage carbohydrates in thiolation mutants. (A) Schematic representation depicting nucleotide and storage carbohydrates (trehalose and glycogen) biosynthesis, starting from a common precursor, glucose-6-phosphate. In normal conditions, flux is typically higher towards the pentose phosphate pathway and eventually nucleotides. (B) Nucleotide synthesis is decreased in tRNA thiolation mutants. WT and tRNA thiolation mutant cells (*ncs2*Δ) grown in minimal media were pulse-labelled with [U-^13^C_6_]-labelled glucose for 10 minutes to measure newly synthesized nucleotides (GMP and AMP) using mass spectrometry. Percent label incorporation in WT and tRNA thiolation mutant cells was plotted. The incorporation of ^13^C atoms from [U-^13^C_6_]-labelled glucose into nucleotides is represented as M+n, where n is the number of ^13^C-labelled atoms (with all five carbons labelled in the ribose sugar). Data are displayed as means ± SD, n=3 for GMP and n=2 for AMP. *p<0.05, **p<0.01. Also see Figures S3A and S3B. (C) Steady-state trehalose and glycogen amounts are increased in tRNA thiolation mutants. Trehalose and glycogen content of WT and tRNA thiolation mutant cells (*uba4*Δ and *ncs2*Δ) grown in minimal media was plotted. Data are displayed as means ± SD, n=4 biological replicates with two technical replicates each for trehalose, and n=3 biological replicates with two technical replicates each for glycogen. ***p<0.001, ****p<0.0001. Also see Figures S3C. (D) Trehalose synthesis is increased in tRNA thiolation mutant. WT and tRNA thiolation mutant cells (*ncs2*Δ) grown in minimal media were pulse-labelled with [U-^13^C_6_]-labelled glucose for 5 minutes to measure newly synthesized trehalose using targeted LC-MS/MS. Normalized peak area of trehalose in WT and tRNA thiolation mutant cells was plotted. The incorporation of ^13^C atoms from [U-^13^C_6_]-labelled glucose into trehalose is represented as M+n, where n is the number of ^13^C-labelled atoms. Data are displayed as means ± SD, n=3. *p<0.05, **p<0.01.

Next, we directly assessed trehalose and glycogen levels biochemically, as the end-point metabolic readouts of the alternative arm where glucose-6-phosphate is diverted away from the PPP. We first estimated the steady-state amounts of trehalose and glycogen with a biochemical assay, and observed a marked increase in these metabolites in the thiolation mutants (Figure 2C and S3C). Subsequently, we directly estimated flux towards trehalose synthesis, using [U-^13^C_6_]-labelled glucose in a similar metabolic flux measurement as described earlier. Here, we observed a strong increase in the synthesis of trehalose (M+6 and M+12 mass isotopomers) in the tRNA thiolation mutants (Figure 2D). These data together reveal that carbon flux is re-routed in the thiolation mutants towards the storage carbohydrates, and away from nucleotide synthesis. Collectively, these results show that cells lacking tRNA thiolation rewire metabolic outputs towards storage carbohydrates, suggesting a ‘starvation-like’ metabolic state. Notably, this occurs despite the absence of glucose (carbon) or amino acid (nitrogen) limitation in these cells.

### tRNA thiolation couples cellular metabolic state with normal cell cycle progression

Dissecting physiological roles of such a fine-tuning of metabolic outputs can be challenging, and this has been the case for tRNA thiolation mutants. However, a simple yeast system, termed ‘yeast metabolic cycles’ or metabolic oscillations, has been effective in identifying regulators that couple metabolism with cell growth/cell-division (Tu *et al.*, 2005; Slavov and Botstein, 2011). In continuous, glucose-limited cultures, yeast cells exhibit robust metabolic oscillations, which are tightly coupled to the cell division cycle, and where DNA replication and cell division are restricted to a single temporal phase (Tu *et al.*, 2005; Chen *et al.*, 2007). In an earlier study we had observed that tRNA thiolation mutants exhibit abnormal metabolic cycles (Laxman *et al.*, 2013). This was reminiscent of phenotypes exhibited by mutants of cell division cycle regulators (Chen *et al.*, 2007). We therefore asked if tRNA thiolation coupled metabolic and cell division cycles. To test this, we sampled the cells at regular intervals of time during the metabolic cycles, and determined their budding index. While WT cells showed synchronized cell cycle progression, tRNA thiolation mutants showed asynchronous cell division (Figure 3A), suggesting a de-coupling of metabolic and cell division cycles. Given our earlier data showing a metabolic rewiring away from nucleotide synthesis in thiolation mutants, we hypothesized that tRNA thiolation controlled normal cell cycle progression by regulating the balance between nucleotide synthesis, and storage carbohydrate synthesis.

**Figure 3.**
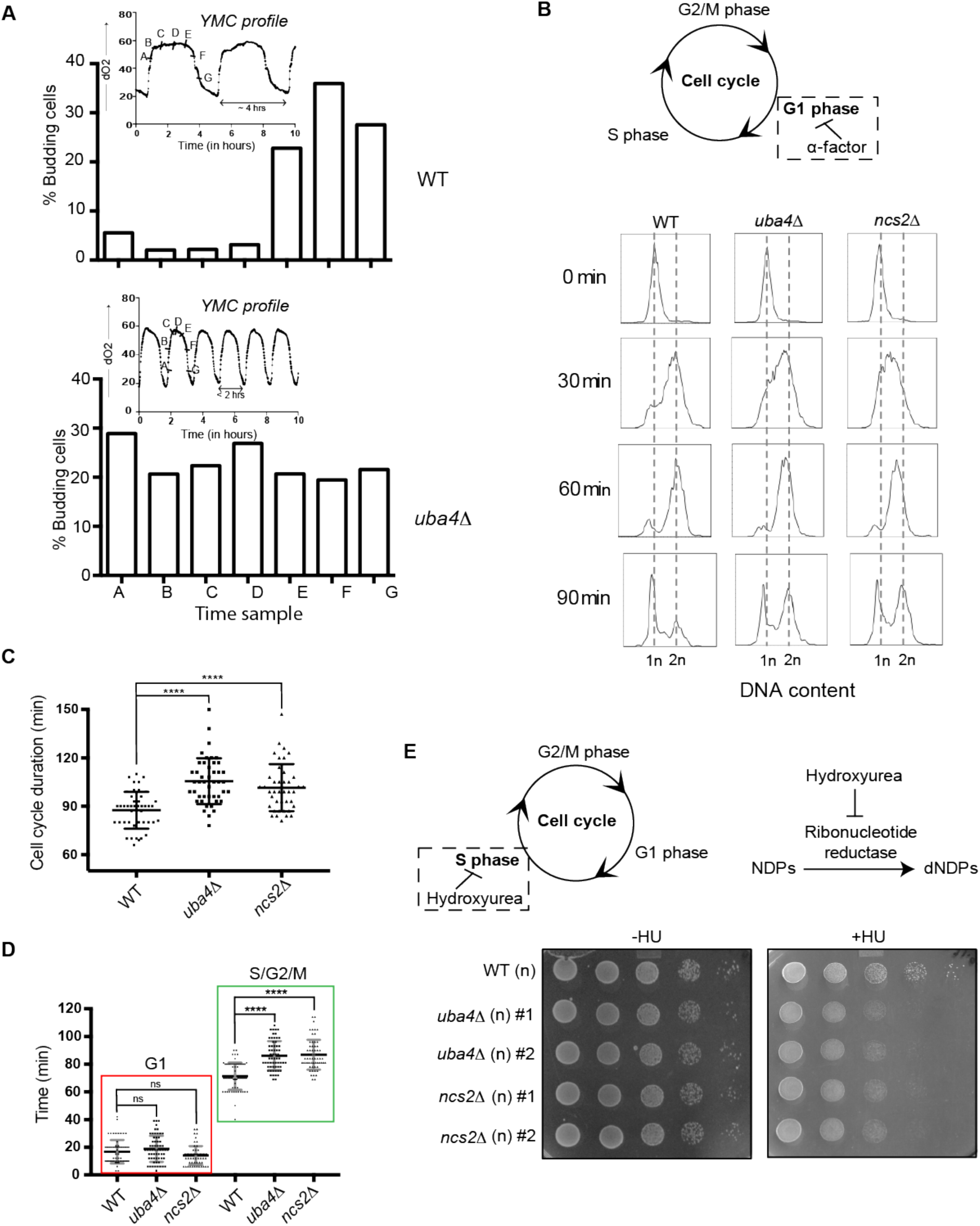
tRNA thiolation couples cellular metabolic state with normal cell cycle progression. (A) tRNA thiolation mutants exhibit asynchronous cell division and disrupted yeast metabolic cycles. WT and tRNA thiolation mutant cells (*uba4*Δ) growing in chemostat cultures under conditions of normal yeast metabolic cycles, were sampled at 20 minute time intervals. Percentage of budded cells represents the fraction of cells in S/G2/M phases of the cell cycle. At least 200 cells were analysed for each time point, for the respective strains. Inset: the oxygen consumption profiles of the WT and tRNA thiolation mutant metabolic cycles are represented. Note that the metabolic cycles of thiolation mutants are disrupted. (B) tRNA thiolation mutants show delayed cell cycle progression. The DNA content of WT and tRNA thiolation mutant cells (*uba4*Δ and *ncs2*Δ) grown in minimal media, during G1 arrest (0 min) and after release from G1-arrest (30, 60 and 90 min) was determined by flow cytometry analysis, and is presented. (C and D) Cell cycle duration, distribution of G1 phase duration (red) and S/G2/M phase durations (green) for WT and tRNA thiolation mutant cells (*uba4*Δ and *ncs2*Δ) were measured in single cells using time lapse live-cell microscopy. We determined the G1 duration (starting from the time of complete division of mother and daughter cells to bud emergence), and S/G2/M durations (from the time of bud emergence to complete division of mother and daughter cells) for only first generation mother cells. At least 100 cells were analysed for each strain. ns denotes non-significant difference. ****p<0.0001. (E) tRNA thiolation mutants exhibit increased HU sensitivity. WT and tRNA thiolation mutant cells (*uba4*Δ and *ncs2*Δ) grown in minimal media were spotted on minimal media agar plates containing 150 mM HU. Also see Figures S4A and S4B.

To test this directly, we arrested cells in G1-phase using alpha factor, synchronously released them into the cell cycle by washing away the alpha factor, and monitored cell cycle progression by flow cytometry (Figure 3B). 30 min post-release from G1-arrest, we observed delayed cell cycle progression and accumulation of cells in the S-phase in tRNA thiolation deficient cells. Further, using time lapse live-cell microscopy we found that the duration of the S-G2/M phase was longer in thiolation mutants (Figure 3C and Figure 3D). Since nucleotides are required for DNA replication during the S-phase of the cell cycle, we reasoned that that this S-phase delay is due to decreased flux towards nucleotide synthesis in thiolation mutants. To investigate this, we examined the sensitivity of WT and thiolation deficient cells to hydroxyurea (HU), which inhibits the ribonucleotide reductase (RNR) enzyme and arrests cells in the S-phase. Thiolation mutants exhibited increased sensitivity to HU (Figure 3E and S4A). This observed HU sensitivity was not due to a defect in the activation of the Rad53 checkpoint pathway in thiolation mutants (Figure S4B).

Summarizing, these results show that tRNA thiolation-mediated regulation of metabolic homeostasis, leading towards regulated nucleotide synthesis, is required for appropriately coupling metabolic state with normal cell cycle progression.

### Loss of tRNA thiolation results in reduced phosphate homeostasis (*PHO*) related transcripts and ribosome-footprints

Thus far, it remains unclear why the loss of tRNA thiolation results in this distinct metabolic switch, where carbon and amino acid flux is diverted away from nucleotide synthesis and into storage carbohydrates. This therefore suggests a deeper, non-intuitive regulatory check-point underpinning the overall metabolic rewiring towards a ‘starvation-like’ state in tRNA thiolation mutants. In order to identify what this controlling bottleneck might be, we identified transcriptional and translational changes in thiolation mutants relative to WT by performing RNA-seq and Ribo-seq (Ingolia *et al.*, 2009) based on methods described in (Weinberg *et al.*, 2016; McGlincy and Ingolia, 2017). Ribosome profiling datasets were generated for both distinct thiolation pathway mutants (*uba4*Δ and *ncs2*Δ), in three biological triplicates. Figures S5A and S5B show transcript and ribosome footprint read correlations, as well as read-length distributions. Using these datasets, we compared global gene expression, as well as ribosome footprints of WT cells with the *uba4*Δ and *ncs2*Δ thiolation mutants (Figure 4A). Notably, comparing WT cells with the thiolation mutants (*uba4*Δ and *ncs2*Δ), we find exceptional correlation for transcripts, as well as ribosome footprints (R^2^>0.97, and p<=2.2x10^-16^ for all datasets) (Figure 4A and 4B). These data surprisingly revealed that there are very little gene expression or translation changes observed in the thiolation mutants, with fewer than ∼30 genes up or downregulated at a two-fold change cutoff (arbitrarily used to illustrate the point), compared to WT cells (Figure 4A and 4B). Furthermore, we observe only modest increases in ribosome-densities at codons recognized by thiolated tRNAs – AAA, CAA and GAA in the *uba4*Δ and *ncs2*Δ cells (Figure S5C). Collectively, these extensive analysis show that the loss of tRNA thiolation has minimal effects on translational outputs *in vivo*, and any changes observed in the translation rates are likely driven by changes at the transcriptional level.

**Figure 4.**
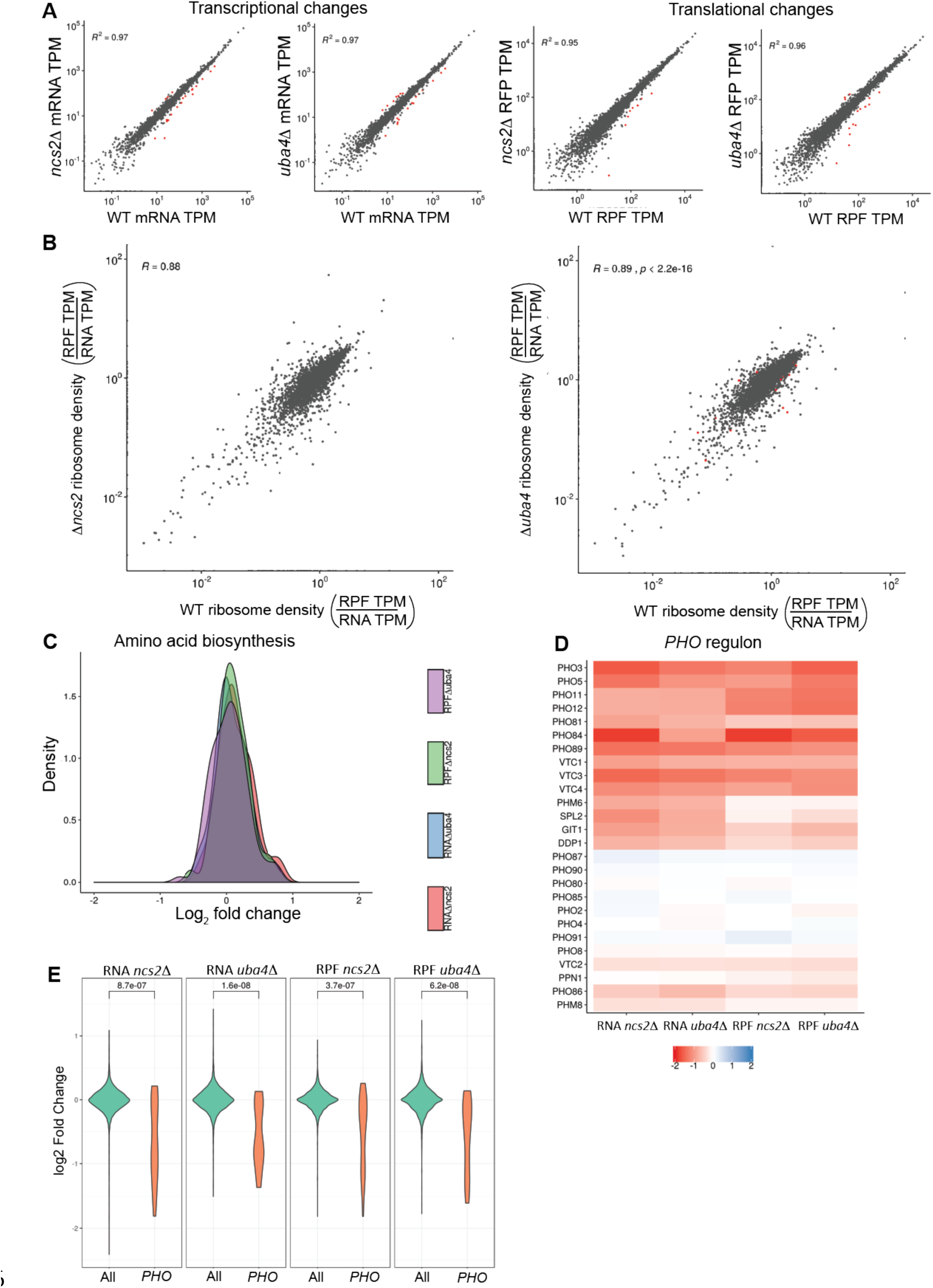
Loss of tRNA thiolation results in reduced phosphate homeostasis (*PHO*) related transcripts and ribosome-footprints. (A) Correlation plots, for WT and tRNA thiolation mutant cells (*ncs2*Δ and *uba4*Δ) for both gene expression (transcript) and ribosome footprint (translation) changes is shown. The coefficients of determination (R^2^) values are shown, along with their significance (p values). Note: Very few differentially regulated genes were observed, and these are indicated as red points. Also see Figures S5A, S5B and S5C. (B) Correlation plots comparing ribosome densities (RPF TPM/RNA TPM) for the thiolation mutants (*ncs2*Δ or *uba4*Δ respectively) with WT cells. Note: overall ribosome densities for thiolation mutants correlate exceptionally well with ribosome densities in WT cells, with the correlation coefficients (R) > 0.88 in both comparisons. (C) A density plot, representing changes in expression (transcript and ribosome footprints) of genes associated with amino acid biosynthetic pathways. In general, amino acid biosynthesis genes were upregulated in tRNA thiolation mutant cells (*uba4*Δ and *ncs2*Δ). Also see Figures S6A, S6B and S6C. (D) Heat map depicting changes in expression (transcript and ribosome footprints) of genes associated with the phosphate (*PHO*) regulon, in the tRNA thiolation mutant cells (*uba4*Δ and *ncs2*Δ), compared to WT cells, at the transcript and ribosome-footprint levels (log2 2-fold changes). (E) A density plot with a statistical comparison between WT and tRNA thiolation mutants, for changes in transcript or ribosome footprints of all genes in the genome, or only the *PHO* regulon. Note: p<10^-7^ for all the comparison datasets. Also see Figure S6D.

Given the lack of large-scale changes at the transcriptional and translational levels, but robust effects of thiolation mutants on cellular metabolism, we focused on changes in expression levels of genes involved in metabolic pathways. We first examined several general amino acid control (GAAC) response genes, including Gcn4 targets in amino acid biosynthetic pathways, and expectedly found these to be transcriptionally upregulated in the thiolation deficient cells (Figure 4C). Additionally, as expected, we observed small increases in *GCN4* translation in the thiolation mutants (Figure S6A). These data collectively corroborate our earlier data from Figure 1, and agrees with previous reports (Zinshteyn and Gilbert, 2013; Nedialkova and Leidel, 2015). Also consistent with our earlier data, most nucleotide biosynthesis genes showed an increase in mRNA and ribosome-footprint abundances in the thiolation deficient cells (Figure S6B). In these datasets, there were also no obvious changes in central carbon metabolism genes in thiolation mutants. We also found small decreases in the transcription and translation rates of all large and small subunit genes of ribosomes in *uba4*Δ and *ncs2*Δ cells at the translational level (Figure S6C), consistent with earlier observations (Laxman *et al.*, 2013; Nedialkova and Leidel, 2015).

However, in the course of this extensive functional, we observed that an unusual group of ∼20 genes were strongly down-regulated in the thiolation mutants, both at the transcript and ribosome-footprint levels (Figure 4D). Although these do not obviously group into a single category based on gene-ontology (GO), we noted that these were functionally related to a metabolic node. These genes are all part of the *PHO* regulon, which regulates phosphate homeostasis in cells (Ljungdahl and Daignan-Fornier, 2012; Secco *et al.*, 2012). This downregulation of these *PHO* related genes was exceptionally significant (p<10^-7^), compared to other genes across the genome, both at the level of transcript abundances and ribosome-footprints (Figure 4E). Also notably, the unaltered gene transcripts/ribosome footprints in the *PHO* regulon (*PHO2, PHO4, PHO80, PHO85, PHO87, PHO90* and *PHO91*) are transcription factors, cyclins/cyclin dependent kinases or low affinity phosphate transporters that are not transcriptionally/translationally regulated, but are regulated at the level of their activity (Lemire *et al.*, 1985; Toh-E and Shimauchi, 1986; Madden *et al.*, 1988; Yoshida *et al.*, 1989; Madden, Johnson and Bergman, 1990; Schneider, Smith and O’Shea, 1994; Ogawa *et al.*, 1995; Lenburg and O’Shea, 1996; Auesukaree *et al.*, 2003). Finally, we also observed that some genes related to phospholipid metabolism were downregulated in the thiolation mutants (Figure S6D). Collectively, these data unexpectedly revealed a strong downregulation of genes related to phosphate homeostasis in the tRNA thiolation mutants.

### Phosphate depletion in wild-type cells phenocopies tRNA thiolation mutants

Inorganic phosphate (Pi) homeostasis is complex, but critical for overall nutrient homeostasis (Ljungdahl and Daignan-Fornier, 2012; Secco *et al.*, 2012). The *PHO* regulon comprises of several genes that respond to phosphate starvation, and maintains internal phosphate levels by balancing transport of Pi from the external environment, from within vacuolar stores, and the nucleus (Figure 5A) (Ljungdahl and Daignan-Fornier, 2012; Secco *et al.*, 2012). Extensive studies have defined general cellular responses to phosphate limitation (Ogawa, DeRisi and Brown, 2000; Wykoff and O’Shea, 2001; Boer *et al.*, 2003, 2010; Saldanha, Brauer and Botstein, 2004; Gresham *et al.*, 2011; Levy *et al.*, 2011; Choi *et al.*, 2017; Gurvich, Leshkowitz and Barkai, 2017). In general, the *PHO* response is very sensitive to phosphate limitation, inducing rapidly to restore internal phosphate levels. Extended phosphate starvation switches cells to an overall metabolically starved state. Our observed reduction in *PHO* related transcripts and ribosome-footprints in the tRNA thiolation mutants was striking. We therefore first biochemically validated our results from the ribosome-profiling data. We measured protein amounts of Pho12 and Pho84, two arbitrarily selected *PHO* genes, in WT cells and the thiolation mutants. These proteins were substantially reduced in *uba4*Δ and *ncs2*Δ cells (Figure 5B). This suggests that the thiolation mutants were effectively in a constitutively phosphate-limited state. To test this, we next asked if phosphate starvation resembles the key metabolic hallmarks of the thiolation mutants. We first biochemically estimated amounts of trehalose in WT cells with or without phosphate starvation, and found a robust increase in trehalose after phosphate starvation (Figure 5C). We next measured Gcn4 (protein) in WT cells, with or without phosphate starvation. Here, we observed a strong induction in Gcn4 protein upon phosphate starvation (Figure 5D). Thus, these data from WT cells starved of phosphate strikingly phenocopied the tRNA thiolation mutants. This is also strikingly consistent with earlier metabolic profiling studies of responses to phosphate limitation, which also observed higher amino acid and low nucleotide levels under these conditions (Boer *et al.*, 2010). We therefore conclude that the tRNA thiolation mutants are effectively phosphate-limited, and have altered phosphate homeostasis, due to a reduction in the *PHO* related genes. This effective phosphate limitation is responsible for the metabolic state switch in these cells.

**Figure 5:**
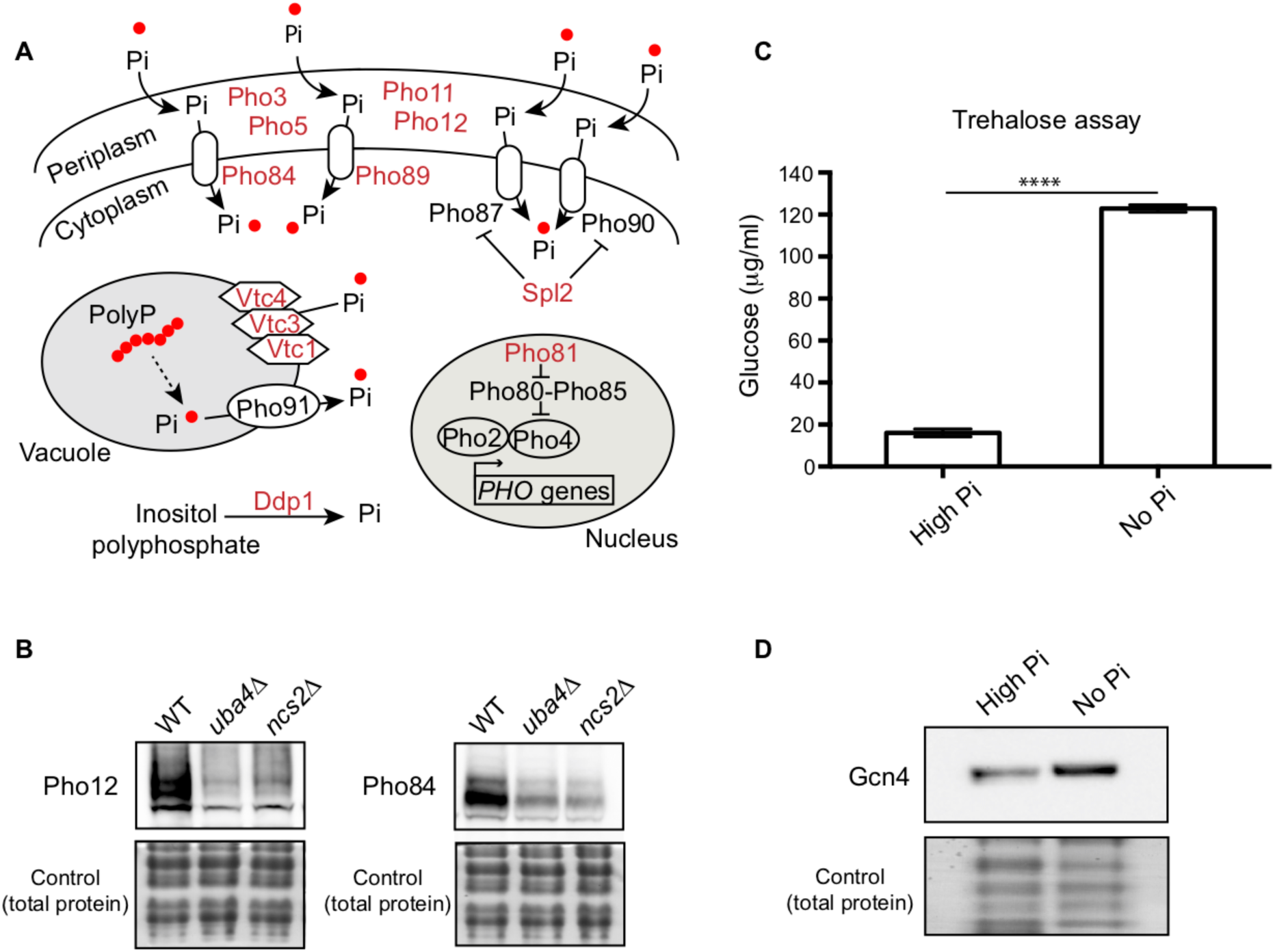
Phosphate depletion in wild-type cells phenocopies tRNA thiolation mutants. (A) A schematic representation of *PHO* regulon related genes, and their roles. Pho84 and Pho89 are high affinity phosphate transporters, Pho87 and Pho90 are low affinity phosphate transporters, Spl2 is a negative regulator of low affinity phosphate transporters, Pho3, Pho5, Pho11 and Pho12 are secreted acid phosphatases, Pho80-Pho85 is a cyclin-dependent kinase (CDK) complex, Pho81 is a CDK inhibitor, Pho2 and Pho4 are transcription factors, Vtc1, Vtc3 and Vtc4 are involved in vacuolar polyphosphate accumulation, Pho91 is a vacuolar phosphate transporter, Ddp1 and Ppn1 are polyphosphatases. Genes marked in red are down-regulated in tRNA thiolation mutant cells (*uba4*Δ and *ncs2*Δ). (B) Pho12 and Pho84 proteins are decreased in tRNA thiolation mutants. Pho12 and Pho84 protein levels (Pho12 and Pho84 tagged with FLAG epitope at their endogenous loci) in WT and tRNA thiolation mutant cells (*uba4*Δ, *ncs2*Δ and *ncs6*Δ) grown in minimal media were detected by Western blot analysis using an anti-FLAG antibody. A representative blot is shown (n=3). (C) Trehalose amounts are increased upon phosphate starvation. Trehalose content of WT cells grown in high and no Pi media was plotted. Data are displayed as means ± SD, n=3. ****p<0.0001. (D) Gcn4 protein is increased upon phosphate starvation. Gcn4 protein levels (Gcn4 tagged with HA epitope at the endogenous locus) in WT grown in high and no Pi media were detected by Western blot analysis using an anti-HA antibody. A representative blot is shown (n=3).

### Trehalose synthesis associated phosphate release enables cells to maintain phosphate balance

This observed downregulation of phosphate metabolism in the thiolation deficient cells is striking. Nonetheless, it is not immediately obvious biochemically how this relates to re-routing carbon towards storage carbohydrates, and decoupling amino acid metabolism from nucleotide synthesis. Perplexingly, in our transcript and translation analysis, no other metabolic arms were similarly decreased in the thiolation deficient cells, and only the amino acid biosynthesis arm (dependent on Gcn4) increases, which we have addressed earlier. Notably, while earlier studies have hinted that phosphate limitation results in a shift towards storage carbohydrates (Lillie and Pringle, 1980; Boer *et al.*, 2003, 2010), this more extensive metabolic rewiring has not been carefully analyzed, and a biochemical explanation for this is missing. We wondered if some overlooked biochemical process could explain why a perturbation in phosphate homeostasis connects to the synthesis of storage carbohydrates trehalose and glycogen, as also seen in tRNA thiolation mutants. To address this, we carefully examined all the metabolic nodes altered in the tRNA thiolation mutants, evaluating necessary co-factors and products of each pathway, and looking for possible connections to phosphate. Here, we noted an apparently minor, largely ignored output in the arm of carbon metabolism, where glucose-6-phosphate is routed towards trehalose synthesis. The first step of trehalose synthesis is the formation trehalose-6-phosphate (T-6-P), carried out by trehalose-6-phosphate synthase (Tps1). This is followed by the dephosphorylation of T-6-P by Tps2 (Virgilio *et al.*, 1993), forming trehalose (Figure 6A). We noted that this Tps2-dependent second step is accompanied by the release of free, inorganic phosphate (Pi) (Figure 6A). Canonically, these two steps are viewed as an apparently futile trehalose cycle during glycolysis, regenerating glucose, in order to maintain balanced glycolytic flux (van Heerden *et al.*, 2014; Van Heerden *et al.*, 2014). However, we reasoned that if the availability of inorganic phosphate is limiting, a shift to trehalose synthesis can be a way by which cells can liberate Pi, and restore phosphate levels. For this to be generally true, the prediction is that during phosphate starvation, WT cells must accumulate trehalose in order to recover phosphate. As shown earlier, this is exactly what is observed in WT cells limited for phosphate (Figure 5C), and in the tRNA thiolation mutants (Figure 2C and 2D) which are effectively phosphate limited due to a reduction in the *PHO* genes.

**Figure 6:**
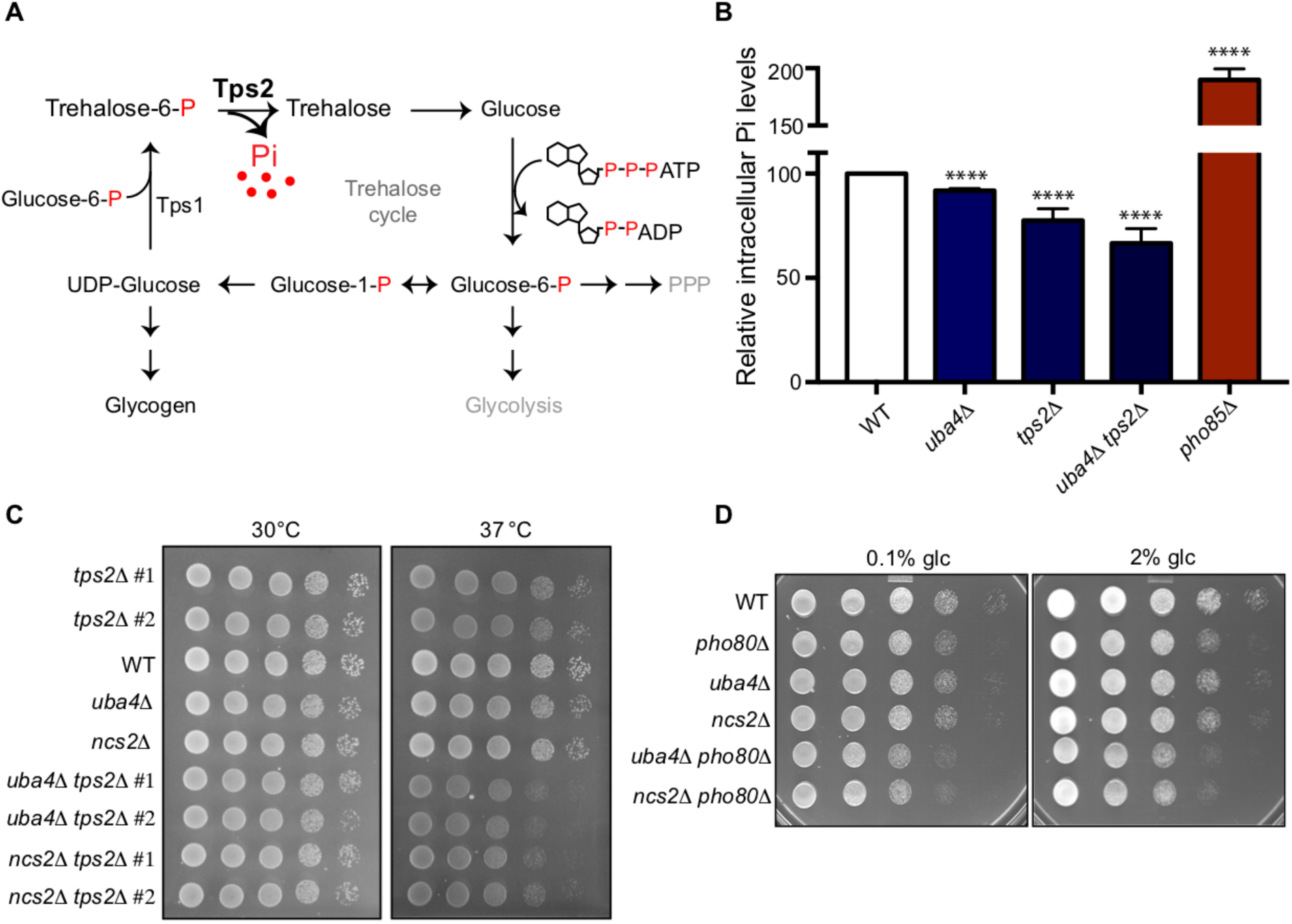
Trehalose synthesis associated phosphate release enables cells to maintain phosphate balance. (A) A schematic representation of the trehalose cycle, showing the routing of glucose-6-phospate towards trehalose and glycogen biosynthesis. Trehalose synthesis requires two glucose molecules, catalyzed by the Tps1 and Tps2 enzymes. The Tps2-mediated reaction synthesizes trehalose, and notably releases free Pi. (B) Intracellular Pi levels are maintained in tRNA thiolation mutant cells (*uba4*Δ) by Tps2 activity. Free intracellular Pi levels were determined in WT, tRNA thiolation mutant (*uba4*Δ), *tps2*Δ, *uba4*Δ *tps2*Δ and *pho85*Δ cells grown in minimal media, by a colorimetric assay. Intracellular Pi levels in mutant cells relative to WT are plotted, where WT was set at 100. Data are displayed as means ± SD, at least n=2 biological replicates with three technical replicates each. ****p<0.0001. Also see Figure S7A. (C) Synthetic genetic interaction between *TPS2* and tRNA thiolation genes (*UBA4* and *NCS2*) at 37º C. WT, tRNA thiolation mutants (*uba4*Δ and *ncs2*Δ), *tps2*Δ, *uba4*Δ *tps2*Δ and *ncs2*Δ *tps2*Δ double mutant cells grown in minimal media were spotted on low Pi media agar plates with 2% glucose and incubated at 30º C and 37º C. Also see Figure S7B. (D) Synthetic genetic interaction between *PHO80* and tRNA thiolation genes (*UBA4* and *NCS2*). WT, tRNA thiolation mutants (*uba4*Δ and *ncs2*Δ), *pho80*Δ, *uba4*Δ *pho80*Δ and *ncs2*Δ *pho80*Δ double mutant cells grown in minimal media (0.1% glucose) were spotted on minimal media agar plates (0.1% and 2% glucose) and incubated at 30º C.

Given the central role of phosphate, cells utilize all means possible to restore internal phosphate (Ljungdahl and Daignan-Fornier, 2012). Therefore it is experimentally challenging to study changes in phosphate homeostasis in cells. However, we directly tested the hypothesis that trehalose synthesis is a direct way for cells to restore internal phosphate in tRNA thiolation mutants, by utilizing cells lacking *TPS2.* These cells cannot complete trehalose synthesis, and importantly cannot release phosphate (Figure 6A). We first measured the intracellular Pi levels in WT cells, thiolation mutants (*uba4*Δ), cells lacking Tps2p (*tps2*Δ), and cells lacking tRNA thiolation as well as Tps2p (*uba4*Δ *tps2*Δ). *pho85*Δ cells were used as a control, since they exhibit intrinsically higher intracellular Pi levels (Liu *et al.*, 2017). We observed that in cells lacking *TPS2* (*tps2*Δ) intracellular Pi levels were substantially lower (∼75-80%) relative to WT cells (Figure 6B and S7A). This suggests that while other pathways (phosphate uptake, glycerol production and vacuolar phosphate export) remain relevant, Tps2p-mediated Pi release by dephosphorylation of trehalose-6-P is itself important for maintaining internal phosphate levels. Importantly, *uba4*Δ cells had only slightly reduced intracellular Pi levels (∼90%) relative to WT cells (Figure 6B and S7A). This is consistent with the prediction that due to reduced *PHO* expression in these cells, they compensate phosphate through increased trehalose synthesis. Contrastingly, the *uba4*Δ *tps2*Δ cells showed a dramatic reduction in Pi levels (∼65%), compared to either of their single mutants. This striking reduction in Pi levels in these cells is consistent with the predicted outcome, where an inability to release phosphate from trehalose (*tps2*Δ) is also coupled with reduced expression of phosphate assimilation genes (*uba4*Δ). Next, we tested possible genetic interactions between *tps2*Δ and thiolation mutants (*uba4*Δ and *ncs2*Δ) by assessing relative growth. In our genetic background, *tps2*Δ cells exhibit slightly slower growth at 37°C. Strikingly, in cells lacking both *TPS2* and tRNA thiolation (*tps2*Δ *uba4*Δ or *tps2*Δ *ncs2*Δ), we observed a strong synthetic growth defect, in conditions of low phosphate as well as normal phosphate (Figure 6C and S7B). This is also consistent with the proposed role of Tps2p in maintaining phosphate balance in thiolation mutants. Finally, if we completely imbalance phosphate homeostasis in cells, using cells lacking *PHO80*, individual mutants of either *pho80*Δ or thiolation deficient cells show minimal growth defects, but double mutants (*pho80*Δ *uba4*Δ or *pho80*Δ *ncs2*Δ) show a severe synthetic growth defect (Figure 6D).

Collectively, our results suggest that altering phosphate homeostasis by decreasing *PHO* activity regulates overall carbon and nitrogen flow. Cells deal with decreased phosphate availability by diverting carbon flux away from nucleotide biosynthesis, and towards Tps2-dependent trehalose synthesis and Pi release. This restores phosphate, while concurrently resulting in an accumulation of amino acids, and a reduction in nucleotide synthesis.

## Discussion

In this study, we highlight two related findings-a direct role for a component of translational machinery, U_34_ thiolated tRNAs, in regulating cellular metabolism by controlling phosphate homeostasis; and a biochemical rationale for how phosphate availability regulates flux through carbon and nitrogen metabolism.

A comprehensive model emerges from our studies, explaining how high amounts of thiolated tRNAs reflect a ‘growth state’, while reduced tRNA thiolation reflect a ‘starvation state’ (Fig 7). Here, the metabolic control point is phosphate homeostasis. In our proposed model, tRNAs are thiolated in tune with methionine and cysteine availability (Laxman *et al.*, 2013). The presence of these sulfur amino acids reflects an amino acid sufficiency state, suitable for growth (Walvekar, Srinivasan, *et al.*, 2018). In these conditions with high amounts of thiolated tRNA, cells thereby direct carbon flux towards nucleotide biosynthesis, coupled with amino acid utilization (as shown in Figs 1 and 2). Accordingly, at this level of metabolic coupling, thiolated tRNAs sense amino acids, and ensure appropriate nucleotide levels for growth and cell cycle progression (as shown in Figs 2 and 3). On the other hand, a reduction in thiolated tRNAs (during methionine/cysteine limitation) (Laxman *et al.*, 2013) switches cells to a ‘starvation-like state’, by rewiring carbon and nitrogen flux away from nucleotide synthesis and towards storage carbohydrates. This is achieved within the cell by down-regulating the *PHO* regulon, effectively limiting phosphate availability (as shown in Figs 4 and 5). In order to restore phosphate, cells divert glucose flux towards Tps2-mediated trehalose synthesis, and concurrently release Pi (as shown Fig 5 and 6). Thus, while the trehalose shunt and phosphate recycling restores phosphate levels, this is at the cost of decreased nucleotide biosynthesis, and delayed cell cycle progression. Effectively, the loss of tRNA thiolation rewires cells to a starvation state. Collectively, our study reveals how tRNA thiolation appropriately regulates metabolic outputs by controlling phosphate homeostasis, thereby enabling cells to commit to growth (Fig 7). Intriguingly, this correlation of tRNA thiolation with growth and rewired metabolism is now emerging in cancer development (McMahon and Ruggero, 2018; Rapino *et al.*, 2018), suggesting possibly conserved metabolic roles for these modified tRNAs.

**Figure 7:**
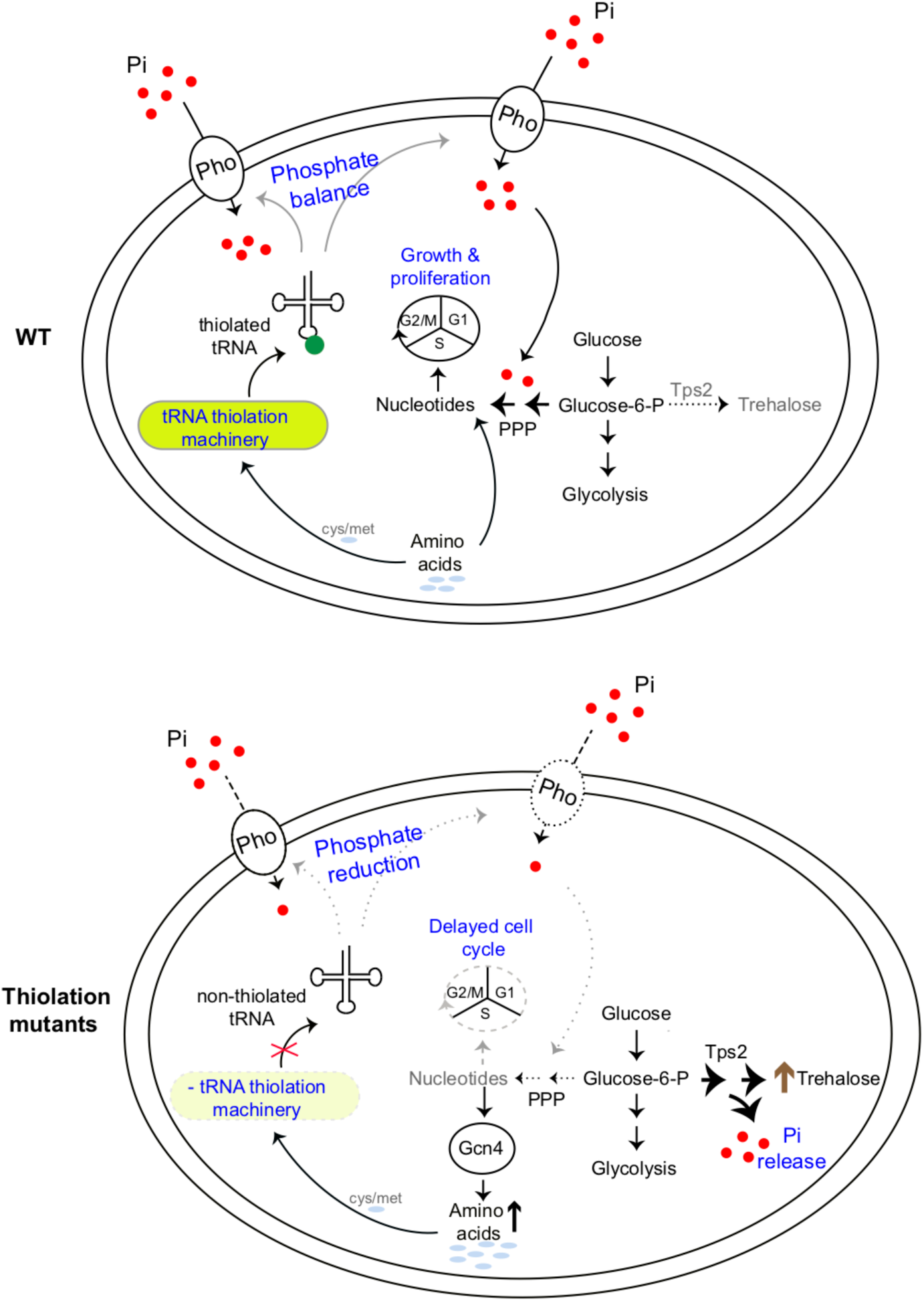
A simple model to illustrate the importance of tRNA thiolation in determining the metabolic state of the cell. WT cells (which have a functional tRNA thiolation machinery) sense and utilize amino acids (methionine and cysteine), and correspondingly have high amounts of thiolated tRNAs. In these cells, carbon flux coupled with amino acid utilization is towards nucleotide biosynthesis. Sufficient nucleotide levels support growth, and proper cell cycle progression, and reflect an overall ‘growth’ metabolic state. In tRNA thiolation mutants, the absence of tRNA thiolation machinery results in non-thiolated tRNAs, and carbon flux is driven away from nucleotide synthesis and towards storage carbohydrates, with a concurrent accumulation of amino acids. This metabolic rewiring in these mutants is due to reduced expression of genes involved in phosphate-responsive signalling pathway, which results in an effectively phosphate-limited state. Due to this, the thiolation mutants attempt to restore intracellular phosphate levels via Tps2-dependent trehalose synthesis, accompanied by phosphate release. Thus, despite the absence of carbon or nitrogen (amino acid) starvation, the tRNA thiolation mutants exhibit an overall metabolic signature of a ‘starved state’, with decreased nucleotide synthesis and delayed cell cycle progression. Pho: phosphate transporters, Pi: inorganic phosphate, PPP: Pentose Phosphate Pathway, cys: cysteine, met: methionine.

A modified tRNA is an unusual but effective mechanism to coordinately regulate metabolic homeostasis. Notably, while several previous studies have observed decreased phosphate-related transcripts in tRNA thiolation deficient cells (Leidel *et al.*, 2009; Nedialkova and Leidel, 2015; Chou *et al.*, 2017), any possible role for phosphate in tRNA thiolation mediated function has been ignored. Furthermore, tRNAs undergo four conserved modifications in the U_34_ position: 5-methoxycarbonylmethyluridine (mcm^5^ U_34_), 2-thiouridine (s^2^ U_34_), 5-methoxycarbonylmethyl-2-thiouridine (mcm^5^s^2^ U_34_), 5-methylaminomethyluridine (mnm^5^U_34_) (refs). Here we focus only on how the s^2^ U_34_ modification (which is derived from sulfur amino acids) regulates cellular metabolic state. Interestingly, the other U_34_ modifications all require s-adenosyl methionine (SAM), and SAM is directly derived from sulfur amino acid metabolism (Thomas and Surdin-Kerjan, 1997). Also interestingly, mutants of all these related U_34_ tRNA modifications show similar metabolic phenotypes as the thiolation mutants (Zinshteyn and Gilbert, 2013; Nedialkova and Leidel, 2015; Chou *et al.*, 2017; Han *et al.*, 2018), and have altered phosphate homeostasis (Chou *et al.*, 2017), raising the possibility that these U_34_-tRNA modifications use similar mechanisms to regulate metabolic homeostasis. A primordial role of U_34_-tRNA modifications might therefore be to sense amino acid sufficiency, and control metabolic state switching towards growth, regulating phosphate availability as a means to achieve this. Co-opting tRNAs (which are the translation components most closely linked to amino acids) to control metabolic states can therefore be an efficient means to ensure appropriate commitments to growth and proliferation, and maximize cellular fitness.

More generally, our findings biochemically explain how phosphate homeostasis determines the extent of carbon and nitrogen flux towards nucleotide synthesis. While it is textbook knowledge that inorganic phosphate is important for glucose homeostasis (Mason *et al.*, 1981; Boyle, 2005; van Heerden *et al.*, 2014; Van Heerden *et al.*, 2014), the biochemical connection of phosphate balance to carbon and nitrogen flux remains poorly explained. Studies have long observed that trehalose increases upon phosphate starvation (Lillie and Pringle, 1980; Klosinska *et al.*, 2011), and *TPS2* is also upregulated (Ogawa, DeRisi and Brown, 2000). In these conditions, central carbon metabolism is down, and phosphate limitation is a ‘general starvation’ cue (Brauer *et al.*, 2008; Boer *et al.*, 2010; Gurvich, Leshkowitz and Barkai, 2017). Why this occurs has not been immediately apparent. Our study, identifying the trehalose shunt as a way to restore phosphate balance, explains these observations. These data also biochemically explain earlier observations from pathogenic fungi, which show that that the amount of trehalose synthesis determines flux through the pentose phosphate pathway, and nitrogen metabolism (Wilson *et al.*, 2007). Additionally, a recent study observed that phosphate starvation results in not just better cell survival in limited nutrient conditions, but also promotes efficient recovery when the nutrients become available (Gurvich, Leshkowitz and Barkai, 2017). Since trehalose accumulation and utilization is tightly coupled with exit and re-entry into cell division cycle respectively (Shi *et al.*, 2010; Shi and Tu, 2013), we propose that trehalose synthesis during phosphate-limitation has dual roles. Trehalose synthesis concurrently releases inorganic phosphate, which restores phosphate balance in the cell and diverts flux away from nucleotide synthesis and growth. When phosphate is no longer limiting, cells can liquidate trehalose to re-enter the cell division cycle, enabling rapid recovery.

Concluding, here we discover that a tRNA modification (thiolated U_34_) enables cells to appropriately balance amino acid and nucleotide levels and regulate metabolic state, by controlling phosphate homeostasis. More generally, we biochemically explain how phosphate homeostasis determines flux through different arms of carbon and nitrogen metabolism.

## Experimental Procedures

### Yeast strains, media and growth conditions

The prototrophic CEN.PK strain of *Saccharomyces cerevisiae* was used in all the experiments (Van Dijken *et al.*, 2000). All the strains used in this study are listed in Table S1. For all experiments, cells were grown overnight at 30ºC in rich media (1% yeast extract, 2% peptone, 2% dextrose), washed once and subsequently sub-cultured in minimal media (0.67% yeast nitrogen base without amino acids, 0.1% glucose) unless specified. Phosphate-limited media was prepared as described previously (Klosinska *et al.*, 2011) except that 0.1%. glucose was used unless specified. The only source of phosphorus in phosphate-limited media (low Pi) was KH_2_PO_4_, which was present at a concentration of 0.15mM. In high phosphate media (high Pi), KH_2_PO_4_ was present at a concentration of 7.5mM with 0.1%. glucose unless specified. In no phosphate media (no Pi), KH_2_PO_4_ was completely absent and 0.1%. glucose was used.

### Western blot analysis

For Gcn4, Pho12 and Pho84 protein levels, cells were grown overnight in rich media, washed once and subsequently sub-cultured in minimal media at an initial OD_600_ of 0.1 and grown till the OD_600_ reaches 0.8-1.0. For Gcn4 protein levels in high and no Pi media, cells were grown overnight in rich media, washed once and subsequently sub-cultured in either high Pi or no Pi media at an initial OD_600_ of 0.1 and incubated for 8 hours at 30ºC. Cells were harvested by centrifugation and protein was isolated by trichloroacetic acid (TCA) precipitation method. Briefly, cells were resuspended in 400 μl of 10% trichloroacetic acid and lysed by bead-beating three times. The precipitates were collected by centrifugation, resuspended in 400 μl of SDS-glycerol buffer (7.3% SDS, 29.1% glycerol and 83.3 mM Tris base) and heated at 100°C for 10 min. The lysate was cleared by centrifugation and protein concentration was determined by using a bicinchoninic acid assay (23225, Thermo Fisher). Equal amounts of samples were electrophoretically resolved on 4-12% pre-cast Bis-tris polyacrylamide gels (NP0322BOX, Invitrogen). Anti-HA (12CA5, Roche) and anti-FLAG (F1804-5MG, Sigma-Aldrich) antibodies were used to detect Gcn4-HA, Pho12-FLAG and Pho84-FLAG proteins. Horseradish peroxidase-conjugated secondary antibodies (mouse and rabbit) were obtained from Sigma-Aldrich. For Western blotting, standard enhanced chemiluminescence reagent (GE Healthcare) was used. Coomassie brilliant blue R-250 was used to stain gels for loading control.

### Metabolite extraction and LC-MS/MS analysis

For steady state amino acids levels, cells were grown overnight in rich media, washed once and subsequently sub-cultured in minimal media at an initial OD_600_ of 0.1 and grown till the OD_600_ reaches 0.8-1.0. ∼10 OD_600_ cells were quenched with 60% methanol at -40°C, and metabolites were extracted, as explained in detail (Walvekar, Rashida, *et al.*, 2018). For ^15^N-label incorporation in amino acids and nucleotides, cells were grown overnight in rich media, washed once and subsequently sub-cultured in minimal media (0.67% yeast nitrogen base without amino acids and ammonium sulfate, 0.1% glucose, 20 mM ammonium sulfate) at an initial OD_600_ of 0.1 and grown till the OD_600_ reaches 0.5. ^15^N_2_-ammonium sulfate (299286, Sigma-Aldrich) was added to reach a ratio of 50% unlabeled to 50% fully labeled ammonium sulfate. Metabolites were extracted from ∼6 OD_600_ cells. For ^13^C-label incorporation in nucleotides and other central carbon metabolites, cells were grown overnight in rich media, washed once and subsequently sub-cultured in minimal media at an initial OD_600_ of 0.1 and grown till the OD_600_ reaches 0.5. [U-^13^C_6_] glucose (CLM-1396-PK, Cambridge Isotope Laboratories) was added to reach a ratio of 50% unlabeled to 50% fully labeled glucose. Metabolites were extracted from ∼6 OD_600_ cells. Extensive metabolite extraction protocols are described (Walvekar, Rashida, *et al.*, 2018). Metabolites were analyzed using LC-MS/MS method as described (Walvekar et al, 2018). Standards were used for developing multiple reaction monitoring (MRM) methods on Thermo Scientific TSQ Vantage Triple Stage Quadrupole Mass Spectrometer or Sciex QTRAP 6500. All the parent/product masses relevant to this study are listed in Table S3. Amino acids were detected in the positive polarity mode. For nucleotide measurements, nitrogen base release was monitored in the positive polarity mode. Trehalose was detected in the negative polarity mode. For PPP metabolites and other triose phosphates, phosphate release was monitored in the negative polarity mode.

Metabolites were separated using a Synergi 4µ Fusion-RP 80A column (100 × 4.6 mm, Phenomenex) on Agilent’s 1290 infinity series UHPLC system coupled to the mass spectrometer. For positive polarity mode, buffers used for separation were-buffer A: 99.9% H_2_O/0.1% formic acid and buffer B: 99.9% methanol/0.1% formic acid (Column temperature, 40°C; Flow rate, 0.4 ml/min; T = 0 min, 0% B; T = 3 min, 5% B; T = 10 min, 60% B; T = 11 min, 95% B; T = 14 min, 95% B; T = 15 min, 5% B; T = 16 min, 0% B; T = 21 min, stop). For negative polarity mode, buffers used for separation were-buffer A: 5 mM ammonium acetate in H_2_O and buffer B: 100% acetonitrile (Column temperature, 25°C; Flow rate: 0.4 ml/min; T = 0 min, 0% B; T = 3 min, 5% B; T = 10 min, 60% B; T = 11 min, 95% B; T = 14 min, 95% B; T = 15 min, 5% B; T = 16 min, 0% B; T = 21 min, stop). The area under each peak was calculated using Thermo Xcalibur software (Qual and Quan browsers) and AB SCIEX MultiQuant software 3.0.1.

### Spotting assay for comparative cell growth estimation

For all spotting assays, cells were grown overnight in rich media, washed once and subsequently sub-cultured in minimal media at an initial OD_600_ of 0.2-0.25 and grown till the OD_600_ reaches 0.8-1.0. Cells were harvested by centrifugation, washed once with water and 10 µl sample for each suspension was spotted in serial 10-fold dilutions. For 8-aza adenine and hydroxyurea sensitivity assays, cells were spotted onto minimal media plates containing 250 and 300 μg/ml 8-aza adenine (A0552, TCI chemicals) or 150 mM hydroxyurea (H8627, Sigma-Aldrich) and incubated at 30ºC. For control plates without drug, images were taken after 1-2 days and for drug containing plates after 4–5 days. For genetic interaction analysis with Tps2, cells were spotted onto high and low Pi media plates with 2% glucose. Plates were incubated at 30ºC and 37ºC. For genetic interaction analysis with Pho80, cells were spotted onto minimal media plates with 0.1% and 2% glucose. Plates were incubated at 30ºC.

### Trehalose and glycogen measurements

For trehalose and glycogen measurements in wild-type and thiolation mutants, cells were grown overnight in rich media, washed once and subsequently sub-cultured in minimal media at an initial OD_600_ of 0.1 and grown till the OD_600_ reaches 0.8-1.0. For trehalose measurement in high and no Pi media, cells were grown overnight in rich media, washed once and subsequently sub-cultured in either high or no Pi media at an initial OD_600_ of 0.1 and incubated for 8 hours at 30ºC. Cells were harvested by centrifugation and washed with ice-cold water. Cells were lysed in 0.25 M sodium carbonate by incubating at 95-98°C for 4 hours. Subsequently, added 0.15 ml 1M acetic acid and 0.6 ml of 0.2 M sodium acetate to bring the solution to pH 5.2. Trehalose and glycogen were digested overnight using trehalase (T8778, Sigma-Aldrich) and amyloglucosidase (10115, Sigma-Aldrich) respectively. Glucose released from these digestions was measured using a Glucose (GO) Assay Kit (GAGO20, Sigma-Aldrich). The concentration of released glucose (µg/ml) was determined from the standard curve and plotted. Statistical significance was determined using Student *T*-test (GraphPad Prism 7).

### Phosphate measurement

Cells were grown overnight in rich media, washed once and subsequently sub-cultured in minimal media at an initial OD_600_ of 0.1 and grown till the OD_600_ reaches 0.8-1.0. Free intracellular phosphate levels were determined as described (McNaughton *et al.*, 2010). Briefly, cells were harvested by centrifugation and washed twice with ice-cold water. Cells were lysed by resuspending in 200 µl 0.1% triton X-100 and vortexed for 5 min with glass beads. The lysate was cleared by centrifugation and protein concentration was determined by using bicinchoninic acid assay (23225, Thermo Fisher). 30 µg of whole cell lysate was used for measurement of free intracellular phosphate levels using ammonium molybdate and ascorbic acid colorimetric assay as described (Ames, 1966). Potassium dihydrogen phosphate solution was used for standard curve (0 to 500 µM KH_2_PO_4_). The amount of phosphate was expressed as µM Pi. Statistical significance was determined using a Student *T*-test (GraphPad Prism 7).

### Cell cycle synchronization and flow cytometry analysis

*bar1*Δ::Hyg, *uba4*Δ::NAT *bar1*Δ::Hyg and *ncs*2Δ::NAT *bar1*Δ::Hyg cells were grown overnight in minimal media and subsequently sub-cultured in minimal media at an initial OD_600_ of 0.05 and grown till the OD_600_ reaches 0.2. Cells were harvested by centrifugation, washed with water and resuspended in the same medium containing 10 μg/ml of α-factor (GenScript). Cells were kept at 30°C for 3 hours till complete G1 arrest was observed by light microscopy. Subsequently, 5 ml culture was harvested by centrifugation, washed with water and fixed with 70% ethanol for G1-arrested population. Remaining culture was synchronously released into the cell cycle by washing away the α-factor. Cells were collected at different intervals of time post G1 release, fixed with ethanol, treated with RNaseA (R4875, Sigma-Aldrich) and a protease solution (P6887, Sigma-Aldrich) as described (Haase and Reed, Cell cycle, 2002). Cells were stained with SYTOX green (S7020, Invitrogen) and analyzed on BD FACS Verse flow cytometer.

### Time-lapse live cell microscopy

Cells were grown overnight in rich media, washed once and subsequently sub-cultured in minimal media at an initial OD_600_ of 0.1 and grown till the OD_600_ reaches 0.4-0.5. 1.5% agar pads (50081, Lonza) were prepared containing minimal media. The pad was cut into small pieces after it solidified. 2 µl of the cell suspension was placed on the agar pad, which was inverted and placed in a glass bottom confocal dish (101350, SPL Life Sciences) for imaging. Phase-contrast images were captured after every 3 minutes’ interval for total of 360 mins on ECLIPSE Ti2 inverted microscope (NIKON) and 60X oil-immersion objective. Images were stacked and analyzed using ImageJ software. Statistical significance was determined using a Student *T*-test (GraphPad Prism 7).

### Transcriptome and Ribosome profiling analyses

Cells were grown overnight in rich media and subsequently shifted to minimal media till the OD600 reaches 0.5-0.8. Cells were rapidly harvested by filtration and lysed, as described in detail (Weinberg *et al.*, 2016; McGlincy and Ingolia, 2017). For both transcriptome and ribosome profiling analyses, three biological replicates each for WT and tRNA thiolation mutant cells (*uba4*Δ and *ncs2*Δ) were included. Total RNA and ribosome-protected fragments were isolated from the cell lysates and RNA-seq and ribosome profiling were performed, as described (Weinberg *et al.*, 2016; McGlincy and Ingolia, 2017), with minor modifications. Separate 5’ and 3’ linkers were ligated to the RNA-fragment instead of 3’ linker followed by circularization (Subtelny *et al.*, 2014). 5’ linkers contained 4 random nt unique molecular identifier (UMI) similar to a 5 nt UMI in 3’ linkers. During size-selection, we restricted the footprint lengths to 18-34 nts. Matched RNA-seq libraries were prepared using RNA that was randomly fragmentation by incubating for 14 min at 95C with in 1 mM EDTA, 6 mM Na2CO3, 44 mM NaHCO3, pH 9.3. RNA-seq fragments were restricted to 18-50 nts. Ribosomal rRNA were removed from pooled RNA-seq and footprinting samples using RiboZero (Epicenter MRZH116). cDNA for the pooled libraries were PCR amplified for 16 cycles.

### Ribosome profiling data processing and analysis

RNA-seq and footprinting reads were mapped to the yeast transcriptome using the riboviz pipeline (Carja *et al.*, 2017). Sequencing adapters were trimmed from reads using Cutadapt 1.14 (Martin, 2011) (--trim-n -e 0.2 –minimum-length 24). The reads from different samples were separated based on the barcodes in their 3’ linkers using fastx_barcode_splitter (FASTX toolkit, Hannon lab) with utmost one mismatch allowed. UMI and barcodes were removed from reads in each sample using Cutadapt (--trim-n -m 10 -u 4 -u -10). Trimmed reads that aligned to yeast rRNAs and tRNAs were removed using HISAT2 v2.1.0 (Kim, Langmead and Salzberg, 2015). Remaining reads were mapped to a set of 5,812 genes in the yeast genome (SGD version R64-2-1_20150113) using HISAT2. Only reads that mapped uniquely were used for all downstream analyses. Codes for generating processed fastq and gff files were obtained from riboviz package (https://github.com/shahpr/RiboViz, (Carja *et al.*, 2017). Gene-specific fold-changes in RNA and footprint abundances were estimated using DESeq2 packages in R (Love, Huber and Anders, 2014) using default log-fold-change shrinkage options. Changes in ribosome-densities (translation efficiencies) were estimated using the Riborex package in R (Li *et al.*, 2017).

The complete transcript/ribosome footprint datasets are available at GEO (number GSE124428). Link: https://www.ncbi.nlm.nih.gov/geo/query/acc.cgi?acc=GSE124428

### Continuous chemostat culture growth to study yeast metabolic cycles, and microscopic analysis

Continuous chemostat cultures to establish the YMC were performed as described previously (Tu et al, Science, 2005). An overnight batch culture of prototrophic CEN.PK strain (van Dijken et al, Enzyme Microb Technol, 2000) grown in rich medium was used to inoculate working volume of 1L in the chemostat. At 20-minute time-intervals, cells were fixed with 2% paraformaldehyde, and imaged under a bright-field microscope. ∼200 cells from each time point were sampled, and budding cells were counted manually.

## Author abbreviations

RG and SLa conceived the project. RG, PS and SLa designed the study. RG, ASW, SL and ZR performed experiments. RG and SL prepared RNA-seq and ribosome-profiling libraries. PS performed detailed analysis of RNA-seq and ribosome-profiling data. RG and SL wrote the manuscript, with contributions from the other authors.

Note: SL (Shun Liang), Sla (Sunil Laxman).

## Acknowledgements

We acknowledge the extensive use of the NCBS/inStem/CCAMP mass spectrometry facility for LC-MS/MS instrument support. We also thank Dr. Anjana Badrinarayan for the use of her live-cell imaging microscope system. We thank Claudio de Virgilio, Nikolai Slavov, Sider Penkov, and Sriram Varahan for critical comments on this manuscript. RG and ASW are supported by the Department of Science and Technology and Science and Engineering Research Board (DST-SERB) national postdoctoral fellowships (PDF/2016/000416 and PDF/2015/000225 respectively). SLa is supported by an Intermediate Fellowship from the Wellcome Trust-DBT India Alliance (grant number IA/I/14/2/501523), as well as institutional support from inStem and the Dept. of Biotechnology (Govt. of India). PS is supported by grants NIH R35 GM124976 and start-up funds from the Human Genetics Institute of New Jersey at Rutgers University

